# Biophysical and functional characterization of K^+^-Cl^-^ co-transporters from *Drosophila melanogaster* and *Hydra vulgaris*

**DOI:** 10.1101/2022.05.12.491617

**Authors:** Satoshi Fudo, Marina Verkhovskaya, Coralie Di Scala, Claudio Rivera, Tommi Kajander

## Abstract

The cation-chloride co-transporter (CCC) superfamily includes ion symporters, which co-transport monovalent cations and Cl^-^. CCCs have crucial roles in shaping signalling and neuronal connectivity in the vertebrate brain. K^+^-Cl^-^ co-transporters (KCCs) are a subfamily of CCCs and carry out the symport of K^+^ and Cl^−^ ions across the plasma membrane. The KCC proteins are involved in various physiological processes, such as cell volume regulation, transepithelial ion transport, synapse formation and signal transmission, and blood pressure regulation.

Among KCCs, KCC2 has gained attention because of its unique and crucial functions in the central nervous system neuronal network. Loss of activity of this transporter has been associated with several neurological disorders including schizophrenia, epilepsy, and chronic pain.

On the other hand, only a limited number of studies of KCCs have been published for invertebrates. Among invertebrate proteins, the *Drosophila melanogaster* KCC (*Dm*KCC) has been studied most and suggested critical for neuronal transmission. Also Cnidarian *Hydra vulgaris* has been shown to have a functional KCC (*Hv*KCC). Comparative analyses of these transporters with vertebrate ones and understanding functional and biophysical aspects of them as a model system can help understand the KCC mechanism of ion transport and its regulation and evolution broadly.

In this study, we chose *Dm*KCC and *Hv*KCC as model systems and purified *Dm*KCC and *Hv*KCC from Sf9 insect cells and characterized their biophysical properties with differential scanning fluorimetry and light scattering techniques. We tested their functionality using a fluorescence assay and developed a method to measure recombinant KCC ion transport activity with flame photometry.

## Introduction

Members of the cation-chloride co-transporter (CCC) superfamily are secondary active ion symporters, which co-transport cations (Na^+^ and/or K^+^) and Cl^-^ [1, 2]. CCCs have crucial roles in shaping proper inhibitory signalling and neuronal connectivity in the vertebrate brain [3]. K^+^-Cl^-^ co-transporters (KCCs) are a subfamily of CCCs that carry out the symport of K^+^ and Cl^−^ ions across the plasma membrane. The KCC proteins are involved in various physiological processes, such as cell volume regulation, transepithelial ion transport, neuroendocrine signaling, synapse formation and signal transmission, and blood pressure regulation [1].

Among KCCs, which include KCC1-4 in most vertebrates [4], in particular KCC2 has gained attention because of its unique and crucial functions in the central nervous system neuronal network [5, 6]. The KCC2 protein is known to be responsible for maintaining low intracellular Cl^−^ concentration at the postsynaptic membrane, which is indispensable for inhibitory signaling by the GABA_A_ receptor [7]. It is also involved in the development of dendritic spines on excitatory glutamatergic synapses through the regulatory domains [8]. The involvement of KCC2 both in excitatory and inhibitory synapses indicates its importance as one of the key molecules for proper signal transmission in the brain. Loss of activity of this transporter has been associated with several neurological disorders including schizophrenia [9], epilepsy [10], traumatic brain injury [11], and chronic pain [12]. KCC2 is considered to be one of the drug targets towards treatment of these diseases [13, 14].

KCC proteins consist of 12 transmembrane α-helix (TM) domain and N- and C-terminal cytoplasmic domains (NTD and CTD). The TM domain is the actual ion transport domain, while NTD and CTD are involved in regulation of transport activity for example through phosphorylation/dephosphorylation of several residues in the domains [6, 15]. CTD is also known to act as a binding scaffold to other proteins. Interactions of KCC2 CTD with Protein Associated with Myc (PAM) [16] and PACSIN1 [17] regulate KCC2 expression positively and negatively, respectively, and affect KCC2-mediated ion flux accordingly, while its interactions with 4.1N [8] and β-PIX [18] are critical for proper dendritic spine formation. Recently, cryo-EM structures of different KCCs were solved, showing an inward-open conformation of the TM domain in each structure [19-24]. Those structures also revealed binding sites for K^+^ and Cl^-^ with highly conserved residues coordinated to those ions.

Because of the electroneutral nature of its ion transport, KCC activity has been analyzed by measuring changes in intra-or extracellular ion concentration [6]. Early work on CCC transporter activity was done with flame photometry [25]. More recently, Tl^+^ and ^86^Rb^+^ ions have often been used as substitutes for K^+^ to detect KCC activity. This is done by measuring fluorescence of Tl^+^-sensitive dye loaded into cells [26] or radioactivity of ^86^Rb^+^ transported into cells [27, 28], respectively, while NH_4_^+^ has also been employed as a K^+^ surrogate to measure KCC activity [29]. On the other hand, the measurement of intracellular Cl^-^ concentration in response to KCC activity has been also done with variable methods including gramicidin perforated patch clamp recording [30], single GABA_A_ channel and NMDA channel recording [31], soma-to-dendrite Cl^-^ gradient [32], quinolinium halide-sensitive indicators [33, 34], and genetically encoded Cl^-^-sensitive indicators [35, 36]. Each of these methods to measure KCC ion transport activity has its advantages and limitations, and therefore it will be still of great benefit to develop alternative methods to detect KCC activity.

To date, only a limited number of studies of invertebrate KCCs have been done. Among the invertebrate proteins, *Drosophila melanogaster* KCC (*Dm*KCC) has been studied the most and suggested to be critical for proper neuronal transmission in *Drosophila* [37, 38]. *Dm*KCC is homologous to mammalian KCC2 and can be inhibited by mammalian KCC2-specific inhibitor, VU0463271 [39, 40]. Cnidarian *Hydra vulgaris* is another species which was shown to have a protein mediating Na^+^-independent K^+^(Tl^+^) and Cl^-^ transport (*Hv*KCC) [41]. Although ion transport activity of *Hv*KCC appears to be significantly lower than *Rattus norvegicus* KCC2 (rat-KCC2), comparative functional, biophysical and structural analyses of this and the *Dm*KCC protein with KCCs from other species could be of great use in order to understand the mechanism of ion transport and its regulation by the KCC transporter family.

In this study, we chose *Dm*KCC and *Hv*KCC as model systems and tested their functionality with a ratiometric Cl^-^-sensitive fluorescent probe [42]. We also developed a novel method to follow ion transport activity of KCCs with flame photometry. In addition, we purified *Dm*KCC and *Hv*KCC from Sf9 insect cells and characterized their thermal stability with differential scanning fluorimetry (DSF) and size-exclusion chromatography-coupled multi-angle static laser light scattering (SEC-MALLS).

## Materials and methods

### Plasmid construction

For insect cell expression, cDNAs coding *Drosophila melanogaster* KCC (*Dm*KCC) variant B (NCBI Reference Sequence: NM_166632.2), *Hydra vulgaris* KCC (*Hv*KCC; NCBI Reference Sequence: XP_012555566.1), and C-terminal domain-truncated *Hv*KCC (*Hv*KCC-ΔCTD) were PCR-amplified with primers containing N-terminal 8xHis and FLAG tags and inserted into the pFastBac1 vector, separately. For mammalian cell expression, cDNAs coding *Dm*KCC, *Hv*KCC and rat-KCC2b were PCR-amplified with primers containing N-terminal FLAG tag and inserted into the pcDNA3.1(-) vector, separately. The constructs were confirmed with DNA sequencing.

### Baculovirus generation and protein expression

The pFastBac1 vectors containing *Dm*KCC, *Hv*KCC, or *Hv*KCC-ΔCTD genes were transformed into *E. coli* DH10Bac cells, and the cells were plated on LB agar plates containing 50 μg/ml kanamycin, 10 μg/ml tetracycline, 7 μg/ml gentamycin, 40 μg/ml isopropyl β-D-1-thiogalactopyranoside (IPTG), 100 μg/ml X-Gal and incubated for 24 h at 37 °C. Positive recombinant colonies (white ones) were selected by the blue-white screening method. The white colonies were picked and re-streaked on another plate and incubated for 24 h. Recombination was further confirmed with colony PCR using M13 primers, and the recombinant bacmid DNA was then extracted. Sf9 cells on a 6-well plate were transfected with the bacmid DNA using TransIT LT1 (Mirus Bio LLC). After incubation at 27 °C for 5-6 days, low-titer virus stock was obtained. High-titer virus stock was obtained by infecting Sf9 cells with the low-titer stock and shaking at 27 °C until the cell viability went below 50%.

For the potassium transport activity assay, Sf9 cells at 1.6 × 10^6^ cells/ml were infected with the high-titer virus stock with multiplicity of infection (MOI) of 2. At 60 h post-infection, the cells were diluted with fresh culture medium to 2.0 × 10^6^ cells/ml and used for the assay. For detergent solubilization screening and protein purification, Sf9 cells at 1.5-2.0 × 10^6^ cells/ml were infected with the high-titer virus stock with MOI of 2. At 60 h post-infection, the cells were harvested by centrifugation at 7000 × g for 10 min.

### Potassium transport activity assay

Sf9 cells infected with baculovirus expressing *Dm*KCC and *Hv*KCC-ΔCTD were used. One milliliter of the cell suspension (2.0 × 10^6^ cells/ml) was centrifuged at 500 × g for 3.5 min, and then the cells were resuspended in 1 ml of the assay buffer (200 mM MES/BTP, pH 6.2, 2 mM MgSO_4_). The reaction was initiated by an addition of RbCl or RbNO_3_ as indicated and after incubation for different times at 20 °C, stopped by centrifugation at 4°C. Then the cells were washed with 1 ml of 500 mM ice-cold mannitol and centrifuged at 700 × g for 3.5 min. The cells pellet was resuspended in 2 ml of Li-test solution, and K^+^ content was measured by flame photometry (PFM 234, Instrumentation Laboratory) immediately.

### Detergent solubilization screening

A cell pellet of infected Sf9 cells was resuspended in 10 mM Tris-HCl pH 7.5, containing protease inhibitor cocktail. The cells were homogenized using a dounce homogenizer with 40 strokes. The cell lysate was centrifuged at 6000 × g for 10 min at 4°C to pellet nuclei and unbroken cells. The supernatant was centrifuged at 150,000 × g for 0.5 h at 4 °C. The resulting pellet was resuspended in PBS plus 100 mM NaCl containing protease inhibitor cocktail. This suspension was then split into aliquots, and each stock detergent solution was added at 1:10 v/v ratio to each aliquot so that each sample contained the desired final concentration of each detergent. For a positive control, milliQ was added instead of a detergent solution to an aliquot (total membrane fraction). The divided samples were then rocked at 4 °C for 16 h. The samples were then centrifuged at 150,000 × g for 0.5 h at 4 °C. The resulting supernatants, detergent-soluble fractions, were loaded on an SDS-PAGE gel. For the positive control (total membrane fraction), the centrifugation step was skipped, and the suspension was loaded on the gel. Proteins were subsequently transferred to PVDF membranes, blocked with 5% milk solution and probed with a mouse anti-FLAG antibody followed by a goat anti-mouse IgG-HRP (horseradish peroxidase)-conjugated antibody. The membrane was incubated with ECL Prime Western Blotting Detection Reagent (GE Healthcare), and then protein bands were detected using ChemiDoc XRS+ (Bio-Rad).

### Protein purification

Cell pellets expressing KCC were resuspended in Lysis buffer (10 mM Tris, pH 7.5, 5 mM MgCl_2_, 10 mM KCl, 0.001 mg/ml DNase I, protease inhibitor cocktail) and then centrifuged at 200,000 × g for 20 min. The pellet was resuspended in Lysis buffer, dounce-homogenized, and then centrifuged at 200,000 × g for 20 min. This step one repeated one more time. The pellet was then resuspended in High-salt buffer (10 mM Tris, pH 7.5, 5 mM MgCl_2_, 10 mM KCl, and 1 M NaCl), dounce-homogenized, and centrifuged at 200,000 × g for 20 min. This step was repeated three more times. The resultant pellet was resuspended in Solubilization buffer (50 mM Na phosphate, pH 7.5, 150 mM NaCl, 50 mM KCl, 10% glycerol) and dounce-homogenized, and 10%/2% *n*-dodecyl-β-D-maltoside (DDM) /cholesteryl hemisuccinate (CHS) stock solution was added to the sample so that the final concentration was 1%/0.2% DDM/CHS. The sample was rocked for 16 h at 4 °C and centrifuged at 170,000 × g for 15 min. The supernatant was incubated with Ni-NTA resin for 3 h at 4 °C in the presence of 10 mM imidazole. The resin was loaded onto a column and washed with Wash buffer (50 mM Na phosphate, pH 7.5, 150 mM NaCl, 50 mM KCl, 10% glycerol, 0.02% DDM, 0.004% CHS, 30 mM imidazole). The KCC protein was eluted with Elution buffer (50 mM Na phosphate, pH 7.5, 150 mM NaCl, 50 mM KCl, 10% glycerol, 0.02% DDM, 0.004% CHS, 250 mM imidazole). The eluate was concentrated to ∼0.5 ml using an Amicon Ultra spin concentrator, 100 kDa cutoff (MilliporeSigma) and run through a Superdex 200 10/300 column (GE Healthcare) with Running buffer (20 mM HEPES pH 7.4, 150 mM NaCl, 50 mM KCl, 0.015% DDM, 0.003% CHS, 5% glycerol). Peak fractions were pooled and concentrated to 1.0 mg/ml.

### Oligomerization state analysis

Size-exclusion chromatography-coupled multi-angle static laser light scattering (SEC-MALLS) was used for characterizing oligomerization states of the purified protein. The protein sample at 1.0 mg/ml was run through a Superdex 200 10/300 column (GE Healthcare) with Running buffer (20 mM HEPES pH 7.4, 150 mM NaCl, 50 mM KCl, 0.015% DDM, 0.003% CHS, 5% glycerol) at a flow rate of 0.3 ml/min with an HPLC system (Shimadzu), a MiniDAWN TREOS light scattering detector, and an Optilab rEX refractive index detector (Wyatt Technology Corp.). Data were then analysed with the protein conjugate program in ASTRA 6 software (Wyatt Technology Corp.).

### Culture and transfection of HEK-293 cells

Human embryonic kidney 293 cells (HEK-293) were maintained in Dulbecco’s Modified Eagle’s Medium (DMEM), supplemented with 10% of fetal bovine serum and 50 IU.mL^-1^ penicillin-streptomycin. Cells were transfected with the appropriate pcDNAs using Lipofectamine reagent 2000 (Life Technologies) and used 40-44 hours after transfection. Briefly, the cells were placed in suspension in Opti-MEM media (6 × 10^5^ cells/mL) and mixed with 150 µL of Lipofectamine / DNA complex. Lipofectamine/DNA complex was obtained by incubating 75 µL of Opti-MEM media with 3.5 µL of Lipofectamine reagent 2000 (mix A containing Lipofectamine 2000). The mix of 75 µL of Opti-MEM and 0.75 µg of DNAs encoding constructs of interest (e.g. 0.5 µg of rat-KCC2 as positive controla and mock-KCC (empty vector) as a negative control, *Dm*KCC or *Hv*KCC and 0.25 µg of SuperClomeleon ; mix B containing DNA) was further prepared. Mixes A and B were mixed, incubated for 20 minutes at room temperature and then incubated with freshly prepared HEK cell suspension. The cells were distributed into a 96-well plate and incubated at 37°C, 5% CO_2_. Then after 12 hours, transfection was terminated by substitution of 90% of the Opti-MEM media by fresh DMEM media. Cells were used in the experiments 40-44 hours after transfection.

### Fluorescent chloride assay

To determine the activity of the rat-KCC2, *Dm*KCC and *Hv*KCC, change in the fluorescence emitted by the Cl^-^ sensitive SuperClomeleon probe [42] in response to change in [Cl^-^]_i_concentration were recorded. Cells expressing rat-KCC2, *Dm*KCC, *Hv*KCC or the mock-KCC2 vector were loaded with [Cl^-^]_i_ by incubating them in solution containing high [K^+^]_o_and [Cl^-^]_o_. This solution named “75K^+^ solution” (containing 148 mM Cl^-^ and 140 mM K^+^ (in mM): 140 KCl, 10 HEPES, 20 D-glucose, 2 CaCl_2_, 2 MgCl_2_, pH 7.4, osmolarity adjusted to 300 mOsm by adding Na-gluconate) was diluted in HEPES-buffered (HBS) solution (in mM) (140 NaCl, 2.5 KCl, 10 HEPES, 20 D-glucose, 2.0 CaCl_2_, 2.0 MgCl_2_, pH 7.4, osmolarity 300 mOsm, adjusted using NaCl). After 10 minutes of loading, a free [K^+^]_o_ (K^+^ free solution containing 148 mM Cl^-^ and 0 mM K^+^ (in mM): 140 NaCl, 10 HEPES, 20 D-glucose, 2 CaCl_2_, 2 MgCl_2_, pH 7.4, osmolarity adjusted to 300 mOsm by adding Na-gluconate) was added in the extracellular media. The addition of the K^+^ free solution changes the sum of electrochemical gradients and provokes KCC-dependent extrusion of Cl^-^ and changes of [Cl^-^]_i._ This triggers modification of the SuperClomeleon fluorescence and a decrease of the ratio of CFP/YFP signal (R_CFP/YFP_) that reflects the functionality of the studied KCC. The fluorescence was measured by using a microplate reader (FluostarOptima, BMG Labtech) with two filter sets activated consecutively. The first measurement was performed using filter set detecting Cl^-^ sensitive fluorescence of YFP part of the SuperClomeleon probe (excitation 500 nm, emission 560 nm) sensitive to Cl^-^, and the second measurement was performed using filter set detecting Cl^-^ insensitive fluorescence of CFP part of the SuperClomeleon probe (excitation 450 nm, emission 480 nm) non sensitive to Cl^-^. The intervals between measurements were 90 seconds. Non-transfected cells were used as control for background fluorescence level. Each experiment was made in triplicate. The results were analyzed using Excel and Graphpad Prism.

### Protein structure modelling

The model for *Dm*KCC was generated using the SwissModel server [43]. Several models were generated with various KCC structure templates. A model generated with KCC3 dimer structure (PDB ID: 6m1y) as a template was chosen because KCC3 has the highest sequence identity to *Dm*KCC (60% identity in the modelled parts). The template structure had the bound K^+^ and Cl^-^ ions in the structure, and no other inhibitors, ligands or mutations that could potentially affect the conformation are present, whilst differences are rather small and other templates could have also been used. The GMQE and QMEAN scores for the model were 0.64 and -4.07, respectively.

## Results and discussion

### Protein expression and purification

Both of the *Hv*KCC and *Dm*KCC constructs were expressed by baculoviral expression from *Spodoptera frugiperda* Sf9 cells, with final yields of 0.1-0.2 mg/l and 0.3-0.4 mg/l for *Hv*KCC and *Dm*KCC, respectively. Solubilization screen indicated (**Fig. 1**) that overall, not much difference was seen between tested maltoside detergents for *Dm*KCC, whilst for the *Hydra vulgaris* protein results indicated that the harsher β-octyl glucoside worked less well. Based on the results, DDM was chosen for both proteins as initial detergent for purification. The *Hv*KCC protein purified with DDM in combination with CHS was in the dimeric state (**Fig. 2**), which is thought to be a functional form for mammalian KCCs [3], hence this was used for both proteins, as is commonly required for many membrane proteins from eucaryotic species. Both proteins were purified by Ni-affinity chromatography followed by size exclusion chromatography, and fractions were pooled for the peak representing the presumed dimer (**Fig. 1**). Purity of the pooled fractions were verified by SDS-PAGE, and the samples were taken further to differential scanning fluorimetry (DSF) and SEC-MALLS analyses.

**Figure 1.**
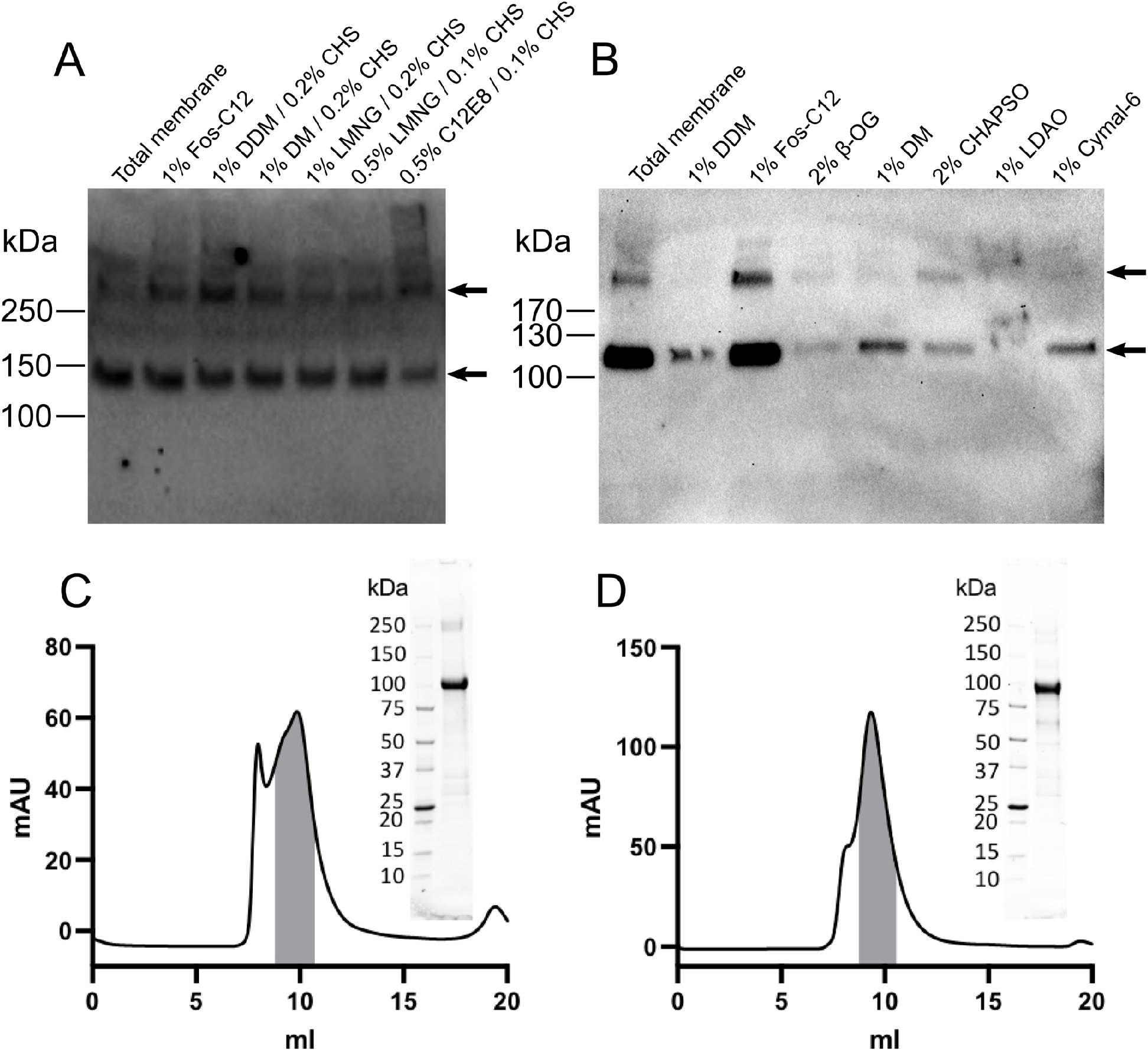
Detergent solubilization screen and purification of KCCs. A) *Dm*KCC and B) *Hv*KCC western blot detection from solubilization with various detergents compared to total membrane preparation. Purification of proteins: C) *Dm*KCC gel filtration profile with collected fractions marked in grey; insert shows the SDS-PAGE of the purified material. D) *Hv*KCC gel filtration profile with collected fractions marked in grey; insert shows the SDS-PAGE of the purified material.

**Figure 2.**
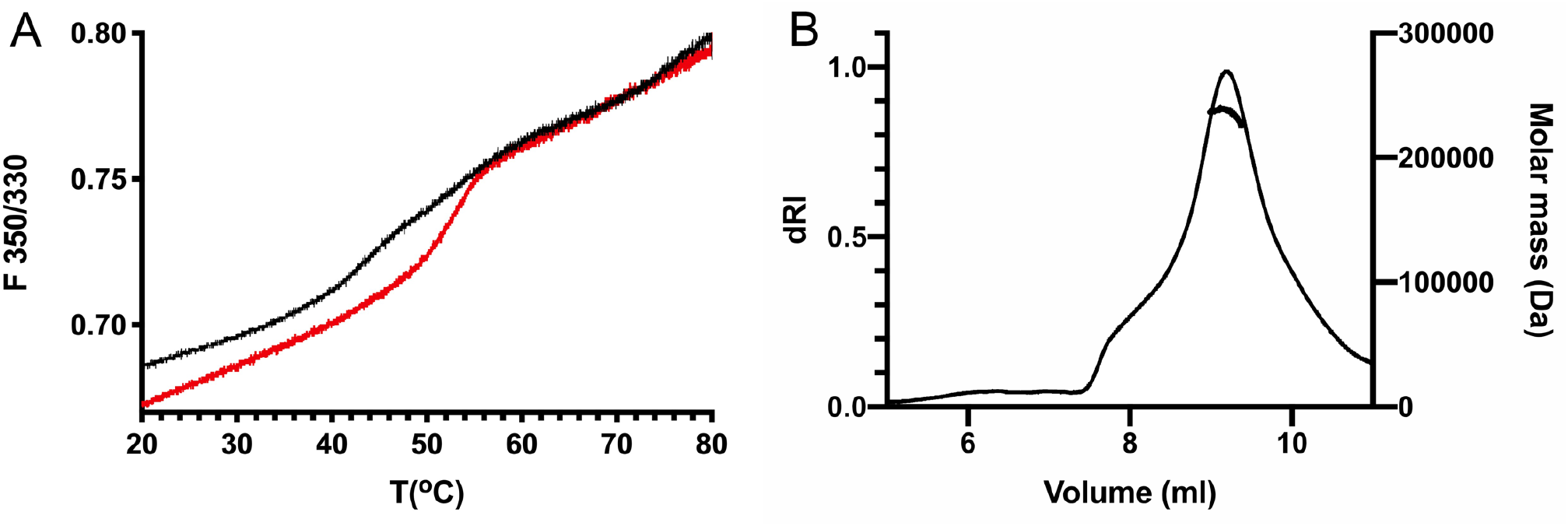
Biophysical characterization of KCCs. A) Differential scanning fluorimetry analysis of the thermostability of *Dm*KCC (red) and *Hv*KCC (black). B) SEC-MALLS analysis of *Hv*KCC shows an apparent dimeric species as the prominent oligomer form with protein molecular mass of ca. 236 kDa, as shown.

### Biophysical characterization

Protein thermostability was analysed by label-free DSF (Prometheus, Nanotemper). *Hv*KCC behaved less well, but *Dm*KCC had a clear transition and was also more stable with T_m_ of approximately 52 °C vs 44 °C for *Hv*KCC as measured from the data (**Fig. 2**).

SEC-MALLS data showed that the *Hydra* protein was mostly dimeric with protein molecular weight of ca. 236 kDa (**Fig. 2**) and total protein:detergent:CHS complex mass of 456 kDa, whilst unfortunately for the *Drosophila* protein, it was not possible to get accurate measurements due to aggregate carryover in light scattering signal. Based on elution profiles (**Fig. 1)** for both proteins, the main peak corresponds to a dimer as verified for *Hv*KCC.

### Activity measurements

As the proteins have not been extensively characterized, we tested in-cell activity of both proteins by the chloride extrusion assay, which shows that the *Drosophila* protein has higher activity, close to the rat-KCC2 control protein, for which the assay has been set up, while the *Hydra* protein was relatively inactive in the HEK-293 cell culture assay (**Fig. 3**). The initial pumping rate is significantly higher for the *Drosophila* protein in this assay, which may mean that there is a difference in functionality of the two proteins or that the *Hydra* protein is differently regulated e.g., by phosphorylation or other mechanisms in the experimental setup in mammalian cell culture.

**Figure 3.**
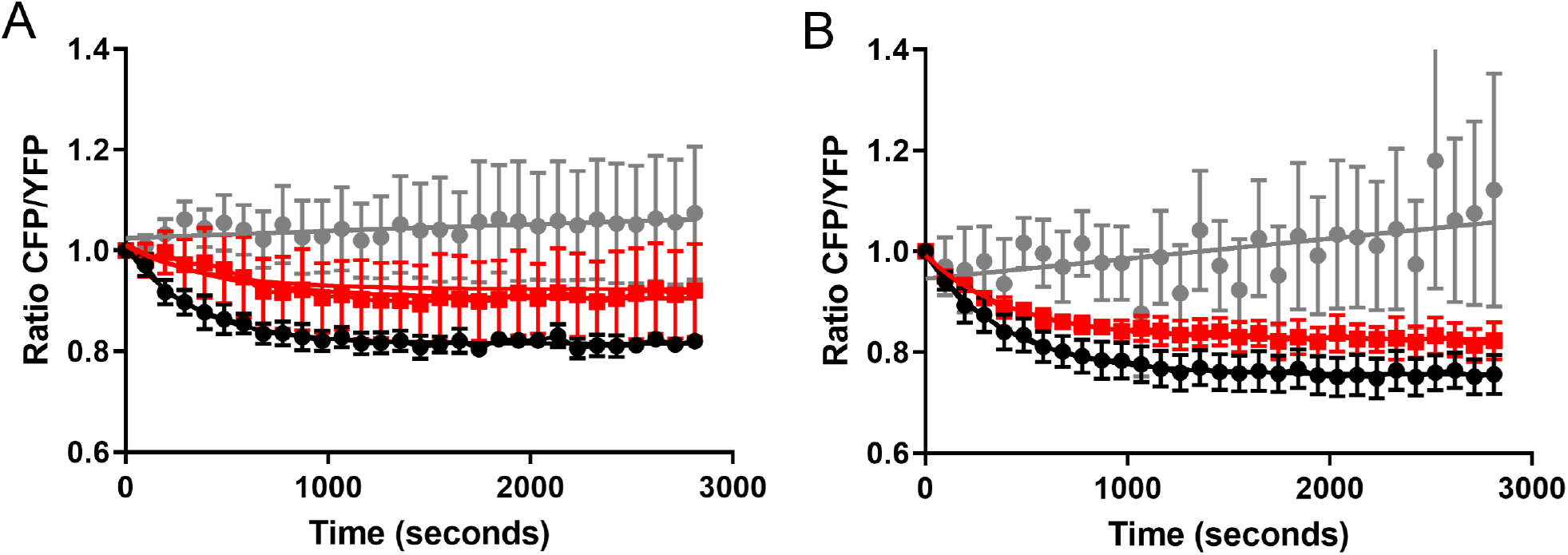
Cell based chloride extrusion assay for KCC activity. Data plotted for A) *Hv*KCC (red squares), B) *Dm*KCC (red squares). The wild type rat-KCC2 used as positive control in A and B (black circles). Cells transfected with the empty vector (grey circles) were used as negative control in both measurements.

Potassium transport activity assays were performed in Sf9 cells to verify activity of the proteins in the expression system (**Fig. 4**). For this purpose, the cells were suspended in K^+^ and Cl^-^ or NO3^-^ free medium and then RbCl or RbNO_3_ were added. Atomic absorption method that discriminates K^+^ and Rb^+^ was used for monitoring K^+^ content in cells. Influx of Rb^+^ resulted in decrease of internal K^+^. Observed K^+^ efflux was strictly dependent on Cl-(but not NO3^-^) and inhibited by bumetanide. This process was associated with active KCC expression, since no such K^+^ efflux was found in Sf9 expressing inactive truncated *Hv*KCC. Therefore, a novel cell-based assay for activity of KCCs was constructed, which can be useful for future studies on ion transporters expressed on insect cells. Activity of *Hv*KCC could not be detected in this assay.

**Figure 4.**
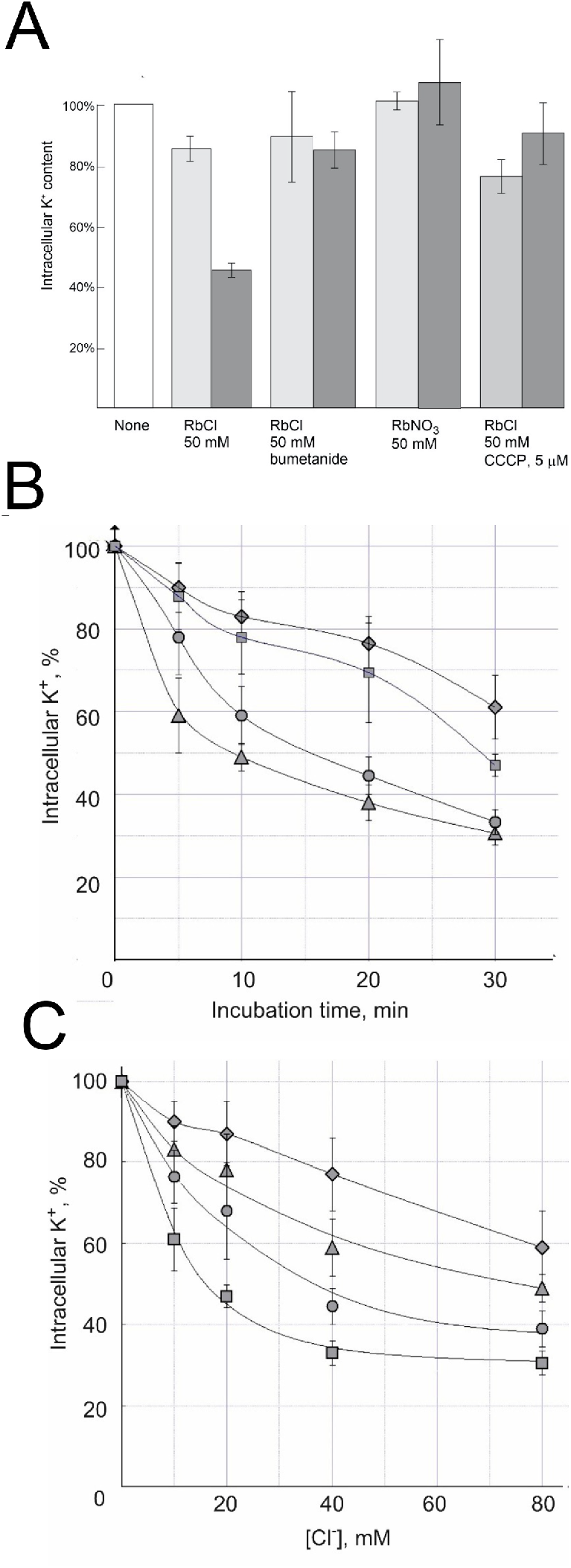
Rb^+^-dependent K^+^ efflux by KCCs from Sf9 cells. A) A bar chart showing *Hv*KCC-ΔCTD (negative control) in light grey and *Dm*KCC in dark grey. B and C) Dependence of Rb^+^-induced K^+^ efflux from Sf9 cells expressing *Dm*KCC on Cl^-^ concentration and the incubation time. B) Dependence of K^+^ efflux on [Cl^-^] at different incubation times: rhombs 5 min, triangles 10 min, circles 20 min, squares 30 min. The solution contained 50 mM RbNO_3_. C) Kinetics of K^+^ efflux at different [Cl^-^]: rhombs 10 mM, squares 20 mM, circles 40 mM, triangles 80 mM. At zero min, 50 mM RbNO_3_ was added.

### Structural and sequence comparison to known KCC structures and implication for ion binding and specificity

Based on the sequence alignment, *Hv*KCC has 51-53% sequence identity to human KCC1-4 and *Dm*KCC, while *Dm*KCC has 55-58% identity to human KCC1-4 and 52 % sequence identity to *Hv*KCC. Based on sequence alignment and structure modelling, the K^+^ and Cl^-^ binding residues in both proteins are conserved similar to human KCC1-4. A model of *Dm*KCC was aligned with human KCC4 structure solved at 2.9 Å resolution [22], which is a highest resolution dimeric wild type structure without inhibitors or mutations and with the ions present. Comparison of the models shows, as expected, that the overall fold is highly similar to the known structures. Both *HvKCC* and *Dm*KCC have the APC family 12-TM helix fold as the other KCCs, and moreover the ion binding residues and their conformation are conserved including residues equivalent to KCC4 K^+^-coordinating (KCC4 numbering) Tyr216 and Thr432 and backbone carbonyls of Asn131 and Thr132, and Cl^-^-coordinating Tyr589 and backbone amides of Val135, Gly433 and Ile444 (**Fig. 5**), while the Na^+^ binding site residues (TM8 Ser614 and Ser165) as seen in NKCC1 structure [44] are not found in *Dm*KCC nor in the *Hv*KCC model or sequence. Thus, the structural modelling as demonstrated in **Fig. 5** for *Dm*KCC shows that the proteins are structurally highly conserved, and the studied proteins have the same functional sites as the mammalian KCCs.

**Figure 5.**
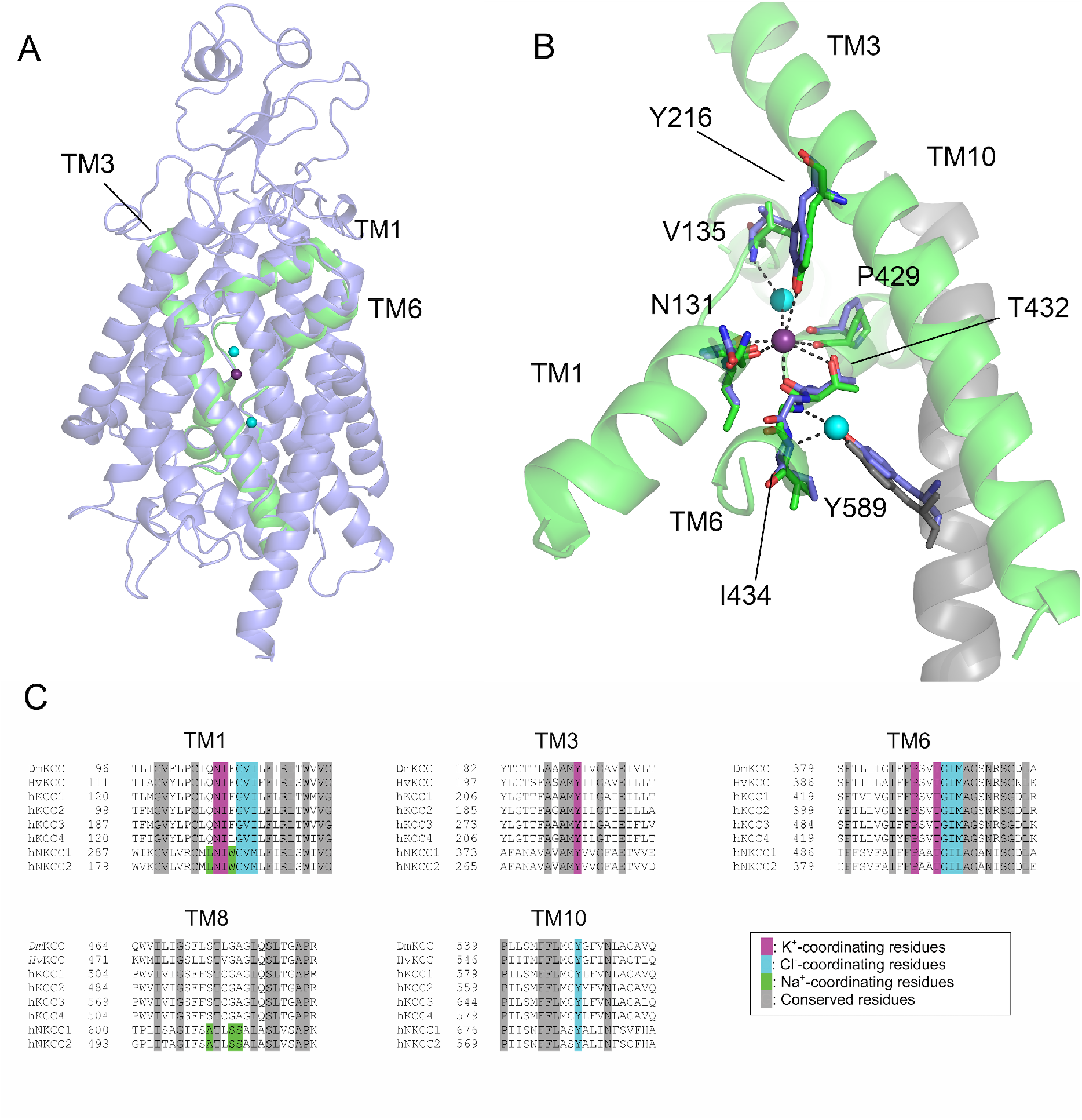
Structural model of *Dm*KCC. A) Overall transmembrane region structure of *Dm*KCC (blue) based on homology modelling compared with KCC4 ion binding helices (in green) with bound Cl^-^ (cyan) and K^+^ (magenta) ions shown as spheres. B) The ion binding site and key residues in TM-helices 1, 3 and 6 (magenta), overlayed with analogous KCC4 residues (green) (KCC4 residue numbering shown, see text). C) Sequence alignment of *Hv*KCC and *Dm*KCC with human KCC1-4 (hKCC1-4) and human NKCC1-2 (hKNCC1-2), with the conserved residues in transmembrane helices TM1, TM3, TM6, TM8 and TM10 coloured as indicated in the figure.

## Conclusions

We have characterized purified KCCs from *Drosophila melanogaster* and *Hydra vulgaris* expressed from insect cell culture and shown that the proteins have K^+^ Cl^-^ transport activity. In particular, purified *Drosophila* protein was shown to be well-behaved in our studies. We also established a cell-based assay to monitor activity of the expressed protein produced for biochemical studies in insect cells, which can be valuable in general for ion transport protein quality control. Detailed characterization of activity of *Hydra* protein was demonstrated earlier [41]. We also showed that the ion binding residues are conserved based on structure prediction from closely related KCC structures recently solved experimentally.

## Acknowledgements

We thank Dr. Maryna Green (University of Helsinki) for help with initial stages of protein expression experiments. The *Hydra* KCC cDNA was a kind gift from Dr. Anna-Maria Hartmann (Carl von Ossietzky University Oldenburg). This research was supported by funding from Jane and Aatos Erkko foundation to TK.

